# Prenatal stress interacts with embryonic loss of *Fgfr2* to increase locomotor hyperactivity in mice

**DOI:** 10.64898/2026.02.04.703738

**Authors:** Michelle X. Chen, Polona Jager, Abigail Sawyer, Hanna E. Stevens

**Affiliations:** Child Study Center, Yale School of Medicine, New Haven, CT, 06520, USA; Department of Psychiatry, Carver College of Medicine, Iowa City, IA, 52242, US

**Author notes:** **Corresponding Author:** Hanna E. Stevens, 1330 Pappajohn Biomedical Discovery Building, 169 Newton Rd, Iowa City, IA 52246.

## Abstract

Attention-deficit/hyperactivity disorder (ADHD) is a developmental psychiatric disorder associated with a complex interplay of genetic and environmental risk factors. We have shown embryonic dorsal forebrain loss in mice of fibroblast growth factor receptor 2 (*Fgfr2*), which has a critical role in normal brain development, results in ADHD-relevant phenotypes: increased locomotion and sociability, and impaired working memory postnatally. How such genetic vulnerabilities interact with environmental exposures to translationally model human ADHD risk remains unclear. Here, we pair the embryonic hGFAP-cre *Fgfr2* conditional knockout (*Fgfr2* cKO) mouse model with prenatal repetitive restraint stress, modeling an environmental factor associated with ADHD risk, to assess adult offspring behaviors and dopamine transporter (DAT) levels. Offspring of prenatally stressed, *Fgfr2* cKO mice show increased locomotion (80% compared to non-stressed, *Fgfr2* cKO animals). Prenatal stress led to a trend increase in impulsivity and trend decrease in working memory but did not affect sociability. There were no interactions with *Fgfr2* cKO observed in these behaviors. Neurobiologically, prenatal stress led to a trend decrease in medial frontal cortex DAT, but these changes did not correlate with behavior. Taken together, our findings implicate prenatal stress as a potential contributor to gene-environment interactions for ADHD risk, supporting its use in translational animal models of childhood psychiatric disorders.

## 1. INTRODUCTION

Attention deficit hyperactivity disorder (ADHD) is a developmental psychiatric disorder, characterized by symptoms of inattention, impulsivity, and/or hyperactivity (1-3). The prevalence of ADHD is currently estimated to be 11.4% of U.S. children having ever been diagnosed (4). Most individuals with ADHD (77.9%) also report experiencing co-occurring disorders, such as anxiety, depression, behavioral conduct difficulties, or learning disabilities (4) which complicate treatment efficacy. As a result, ADHD is a public health concern, underscoring the importance of understanding its etiology to improve prevention and intervention strategies.

As with other brain disorders, ADHD arises from a complex interplay of factors. Twin and adoption studies implicate both genetic and environmental factors in ADHD etiology (1, 2, 5, 6). Despite the relatively high heritability of ADHD, there have been few consistent genetic findings in humans, and the disorder is thought to be polygenic (6-8), making it challenging to behaviorally replicate ADHD with animal models carrying a susceptibility gene. However, emerging transgenic mouse models displaying ADHD-relevant behavioral phenotypes have provided insight into ADHD neurobiological mechanisms even if these target genes do not contribute to polygenic risk. Some models include disruption of fibroblast growth factor (FGF) systems, whose signaling is required for normal formation of the cerebral cortex and affects development of cortical excitatory as well as inhibitory neurons (9-11). Mice with disruptions in FGF receptors (FGFR1, 2, and 3) exhibit a smaller cerebral cortex, which has also been observed in patients with ADHD (12, 13). We recently found embryonic conditional ablation of the fibroblast growth factor receptor 2 (*Fgfr2*) gene in progenitor cells of the dorsal forebrain [(*Fgfr2* conditional knockout (cKO)] (11, 14) in mice leads to behaviors relevant to clinical ADHD presentations, namely hyperactivity, working memory deficits, and social impulsivity in adulthood; interestingly, *Fgfr2* loss in adulthood does not replicate such findings (14). This suggests the embryonic period as a critical neurodevelopmental time for ADHD risk.

One well-documented environmental risk during the embryonic neurodevelopmental period is prenatal stress (15-17), which refers to maternal psychological distress during pregnancy and associated physiological responses (18, 19). Its link to psychopathological outcomes could be explained by its ability to alter intrauterine conditions and thus disrupt normative fetal brain development (15, 20, 21). In human epidemiologic research, prenatal stress frequently associates with ADHD symptoms in the child (16, 22-32). Animal studies further support the link between prenatal stress and ADHD-relevant outcomes in offspring and provide insight into neuronal systems affected by prenatal stress (19, 33-46). However, it is not known to what extent prenatal stress interacts with genetic vulnerability to increase the likelihood of the disorder (26, 47).

Neurobiologically, dysfunction of the dopaminergic (DA) system, particularly in the cortex and the mesolimbic structures, is thought to be implicated in ADHD (7, 48, 49) and in basic functions disrupted in ADHD (50-52). For example, *dopamine transporter* (*DAT*) gene variant (10-repeat SLC6A3 allele) associates with ADHD (53), and *Dat* knockout mice are used to model ADHD (54). Consistent with this, rodent studies of other ADHD risks find alterations in DA systems. For example, prenatal stress in rodents altered *Nurr1* and *Pitx3* expression, which are molecules implicated in DA neuron development, as well as levels of DA, tyrosine hydroxylase (TH), and DA2 receptors (D2R) in the midbrain, nucleus accumbens and cortical regions (19, 55-63). There is some discrepancy, however, in directionality of these changes. Nevertheless, the DA system remains critical to the etiology and treatment of ADHD.

Here, we aim to study a prenatal gene-environment interaction with relevance to ADHD. We leverage the *Fgfr2* cKO model in combination with prenatal stress to assess offspring adult locomotor activity, working memory, social intrusiveness, and impulsivity- and anxiety-relevant behaviors. To start to understand neurobiological mechanisms through which prenatal stress and genetic vulnerability may predispose to ADHD, we assess DAT protein levels, a key player in the DAergic system.

## 2. MATERIALS AND METHODS

### 2.1 Mice

All experimental procedures were approved by the Yale Animal Resources Center and Institutional Animal Care and Use Committee (IACUC) policies in accordance with the National Institutes of Health guidelines. Sufficient animals or samples (n’s for each group noted in each figure caption from 11 non-stressed dams and 8 prenatally-stressed dams) were generated for each assessments using a power analysis based on previous studies and α = 0.05 and β = 0.2.

#### Prenatal restraint stress

Female mice were checked for vaginal plugs daily to detect embryonic day 0. Mice were randomly assigned to prenatal restraint stress (PS) or non-stress, control conditions (NS). Restraint stress was induced in pregnant mice from E12 onwards until birth (P0). The mice were placed in plexiglass restrainers for 45 minutes, three times daily, under bright lights and during the daytime light cycle (42). Control pregnant mice were continuously housed in normal conditions.

#### Generation of Fgfr2 cKO animals

Conditional hGFAP-cre *Fgfr2* knockout mice (referred here as cKO) on a mixed background have been previously described (11, 64). The conditional *Fgfr2* null allele harbors loxP recombination sites flanking regions encoding the Ig III binding and transmembrane domains of the *Fgfr2* gene (*Fgfr2f*) (65). Female mice homozygous for *Fgfr2f* alleles were crossed with male mice homozygous for *Fgfr2f* alleles and also heterozygous for Cre recombinase transgene under the control of the human GFAP (hGFAP) promoter, which is expressed in all dorsal telencephalic radial glia and astrocytes starting from embryonic day (E) 13.5 (66, 67). Cre negative offspring (referred to here as WT), littermates when possible, were used as control animals. Only male offspring are used in this study due to limited resources; childhood psychiatric disorders, particularly ADHD, have higher prevalence in males which allows for these data in males, while limited, to be translationally informative (68).

### 2.2 Behavioral Procedures

All behavioral assessments were performed during the light cycle following at least 30 minutes of habituation prior to testing. Testing order of mice within the cohort was random, each individual mouse was tested on all tasks, and tests were always performed in the same order, as follows, from least to most stressful, as previously (14). In each testing cohort, the mice came from two or more different litters to control for litter effects.

#### Open field

In a chamber (20×14 inches), mice were tested for locomotor activity for 30 minutes each day for two days. Test mice were placed in a chamber corner, and movements were recorded using an overhead camera (AnyMaze videotracking software; Stoelting Co, Wood Dale, Illinois). To control for effects of novelty seeking and/or anxiety on locomotor activity, we present data from the second testing day. The movement velocity and time spent in the center was measured to assess activity level and impulsivity respectively.

#### Y-Maze

In a Y-maze with three arms: A, B, and C (20 cm long, 10 cm wide, 20 cm high, separated by 120°), mice were tested for working memory by being placed in the distal end of arm A and allowed to explore for five minutes. Each individual arm entry (all four paws of the mouse within the arm) and the total number of entries was recorded by an experimenter. The sequence of entries into the maze arms was broken into sets of three consecutive entries, each separated by one entry in the sequence. Spontaneous alternation behavior, defined as the number of sets in which the mouse consecutively entered the three different arms divided by the total number of possible alternations, was used as the measure for working memory (69).

#### Three-chamber social approach

In a three-chamber apparatus, mice were tested for levels of social interaction. Prior to testing, each “stranger” mouse (WT animal of same sex, age, and strain as test mice from a separate cage) was habituated to an inverted wire pencil cup within one side chamber for five minutes. Test mice were habituated to the center chamber for 10 minutes, separated from the other two chambers with doors. Then, the “stranger” mouse (social object) was placed within the inverted cup in one of the side chambers (counterbalanced across test mice), and an identical empty cup (non-social object) in the second side chamber. Movement was recorded using an overhead camera (AnyMaze videotracking software; Stoelting Co, Wood Dale, Illinois) and evaluated for movement around the social vs. non-social objects. Social approach was quantified by comparing the time spent within 1 inch of the social object divided by time spent within 1 inch of the non-social object.

#### Elevated plus maze

For anxiety-relevant behavior assessment, mice were placed at the junction of a maze with four arms, which were elevated from the ground, with two of them closed and two open, to explore the maze for 5 minutes. Movement was recorded using an overhead camera (AnyMaze videotracking software; Stoelting Co, Wood Dale, Illinois) and evaluated for time spent in each arm and center of the maze.

### 2.3 Neuroanatomical measurements

#### Immunohistochemistry

Two weeks after behavioral testing, animals were anesthetized and perfused (phosphate buffered saline (1x PBS), then 4% paraformaldehyde), and brain tissue was post-fixed, cryoprotected (20% sucrose in 1x PBS), embedded in OCT compound, and cryostat (Leica, CM1900, Bannockburn, Illinois) sectioned at 50 μm thickness. Free-floating sections of the medial frontal cortex (mFC, including prelimbic cortex and anterior cingulate cortex), nucleus accumbens (NAc), ventral tegmentum area (VTA), and dorsal striatum were stained using standard immunostaining methods with anti-DAT primary antibody (1:5000, Chemicon) and secondary anti-rat antibody (1:500, Vector Laboratories).

#### Light Microscopy & Punctal Assessment

Digital images were taken of the mFC, VTA, NAc and the striatum using an AxioCam attached to Axio Phot microscope and AxioVision software (Carl Zeiss). Two to three sections (every tenth consecutive section) per region were imaged and analyzed for each brain, bilaterally. The same camera exposure and microscope lighting settings were used each time. Then, DAT puncta were quantified in ImageJ software (National Institutes of Health) by an experimenter blind to the experimental condition of each brain. The DAT immunostaining density values calculated were averaged: first, across sections for a given region for a given brain, and then across the brains in each group for each region separately.

### 2.4 Statistical Analyses

All statistical tests were performed with SPSS version 21. Data normality and variance was assessed for all behavioral and DAT staining density data sets. Any outliers were identified and removed using the modified Thompson tau test (α = 0.05). Outcomes on each behavioral test and for each region’s DAT immunostaining density were normalized to non-stressed (NS), wildtype (WT) littermate controls. Analyses of variance (ANOVA) and ANOVA for repeated measures outcomes were used to assess average velocity and total center time on day 2 in the open field, spontaneous alternation in the Y-maze, social approach in the three-chamber social task, and closed arm time in the elevated plus maze. Pearson correlations were performed to examine associations between DAT density and behavioral measures. Group differences are expressed as percent change relative to control group.

## 3. RESULTS

### 3.1 Prenatal stress may shift ADHD-relevant phenotypes of *Fgfr2* cKO males

To investigate how *Fgfr2 cKO* and prenatal stress independently and interactively impacted ADHD-relevant behaviors, we performed rodent behavioral tasks that allow for the study of modeled ADHD symptoms. Because individuals with ADHD often experience symptoms of hyperactivity, impulsivity, working memory deficits, anxiety, and social functioning challenges, (70) we used four rodent behavioral tasks: open field (to assess locomotor activity and impulsivity-relevant behaviors), Y maze (to assess working memory), three chamber social approach (to assess social intrusiveness), and elevated plus maze (to assess anxiety-relevant behaviors).

#### Hyperactivity

The prenatally stressed *Fgfr2* cKO (PS cKO) group displayed the highest locomotor activity, as measured by average velocity in the open field (Figure 1A): a two-way ANOVA shows prenatal stress interacted with genotype to result in hyperactivity [*F*(1,47) = 16.606, *p* < .0001]. There was a 106% increase relative to the prenatally stressed wild-type (PS WT) offspring, and 80% increase relative to the non-stressed *Fgfr2* cKO (NS cKO) group. No significant change was observed in the PS WT group relative to NS offspring; PS did not exert a main effect [*F*(1,47) = 2.270, *p* = .139]. *Fgfr2* cKO alone increased locomotor activity/hyperactivity [*F*(1,47) = 42.576, *p* < .0001], consistent with our previous study (14). Across-session velocity was plotted (Figure 1B). All groups showed within-session habituation [*F*(5,47) = 40.690, *p* < .0001], but the absolute differences between groups remained the same; all interactions between time factor and other factors were nonsignificant. Finally, the average velocity across the whole field was compared to the less-aversive periphery, and no differences were observed in any of the groups (Figure 1A).

**Figure 1:**
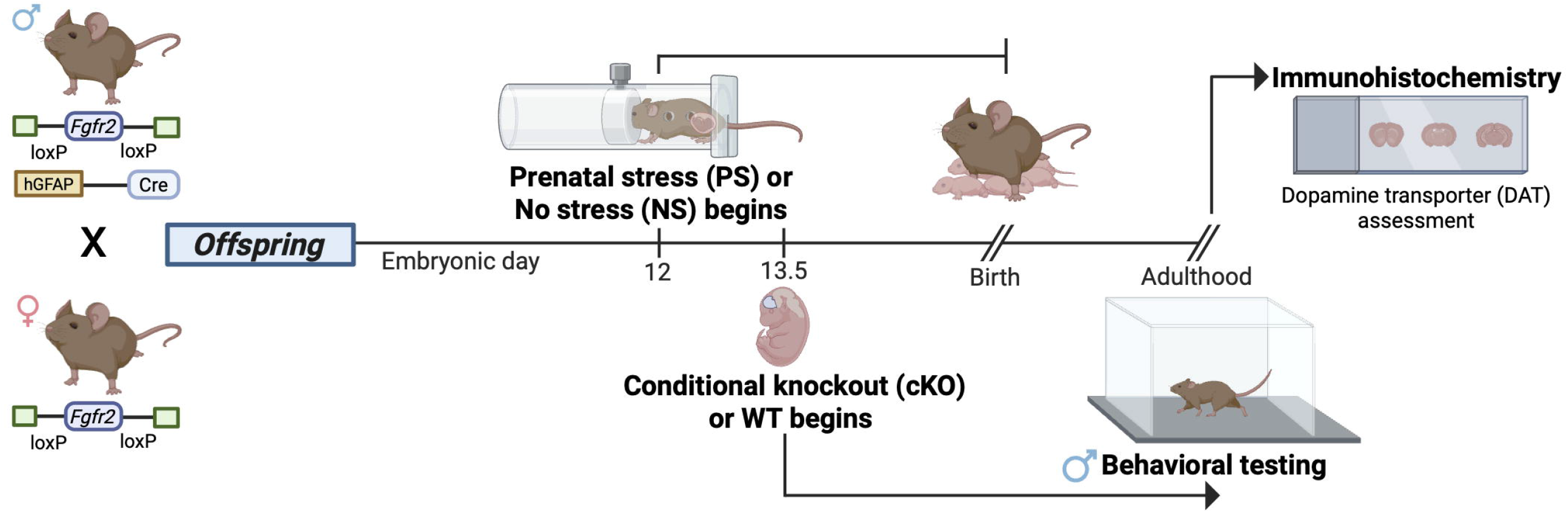
Experimental timeline. Male mice homozygous for Fgfr2f alleles and heterozygous for hGFAP-Cre recombinase were crossed with female mice homozygous for Fgfr2f alleles. Prenatal stress or no-stress conditions began at embryonic day (E)12 until birth. hGFAP expression occurs in all dorsal telencephalic radial glia and astrocytes starting from E13.5. All behavioral testing and neurobiological analyses were performed in adult male mice.

#### Impulsivity

There were no statistically significant differences between groups in their impulsivity, which was measured as center time in the open field on the second day. There was, however, a trend increase in center time in PS offspring [*F*(1,46) = 2.990, *p* = .090], such that PS groups spent on average 19% more time in the center than the non-stressed groups (Figure 1C). The directionality of this effect was in line with the extant literature (44, 45). Given that the whole-field velocity was not lower from the periphery of the field (Figure 1A), this confirms the mice spent time in the center moving.

#### Working memory

There was a trend decrease in working memory performance in PS offspring [*F*(1,48) = 3.154, *p* = .082) and cKO offspring [*F*(1,48) = 2.848, *p* = .098], shown by these trend main effects on spontaneous alternation in the Y-maze (Figure 1D). However, the interaction between the two factors was not significant [*F*(1,48) = .000, *p* = .995), suggesting trend summative deficits. Relative to non-stressed offspring, PS offspring displayed a 10% decrease in spontaneous alternation, and the performance of the cKO animals was also 10% lower compared to the WT mice. All groups performed at or above chance level of spontaneous alternation.

#### Social intrusiveness

Social intrusiveness, as characterized by increased social approach in the three-chamber social task, was not displayed following prenatal stress [*F*(1,47) = 1.554, *p* = .219] (Figure 1E). There was a trend increase in social approach in the cKO groups [*F*(1,47) = 3.371, *p* = .073], the directionality of this effect consistent with the literature on the *Fgfr2* cKO (14). All groups performed above chance level of social approach.

#### Anxiety relevant behavior

There were no main effects of prenatal stress [*F*(1,43) = 1.680, p = .202] or *Fgfr2* cKO [*F*(1,43) = 2.049, *p* = .160] on time spent in closed arms of the elevated plus maze. There was however a trend interaction between prenatal stress and genotype [*F*(1,43) = 3.076, *p* = .087], such that anxiety-relevant behavior of the PS WT animals was increased compared to the rest of the groups. There was an 30% increase in closed arm time relative to the PS cKO and the NS WT groups. This pattern of effects, which is different from those observed for other measures, further suggests impulsivity and hyperactivity in the open field were not confounded by anxiety.

### 3.2 Prenatal stress changed DAT density in the mFC

To explore if abnormalities in DA signaling occurred with behavioral outcomes, we measured dopamine transporter (DAT) protein levels through immunohistochemistry (Figure 2A). DAT uptakes DA into presynaptic terminals and is involved in temporal and spatial control of DA signaling (54). Its expression is tightly regulated (54, 71) and restricted to DAergic cells (72, 73).

**Figure 2:**
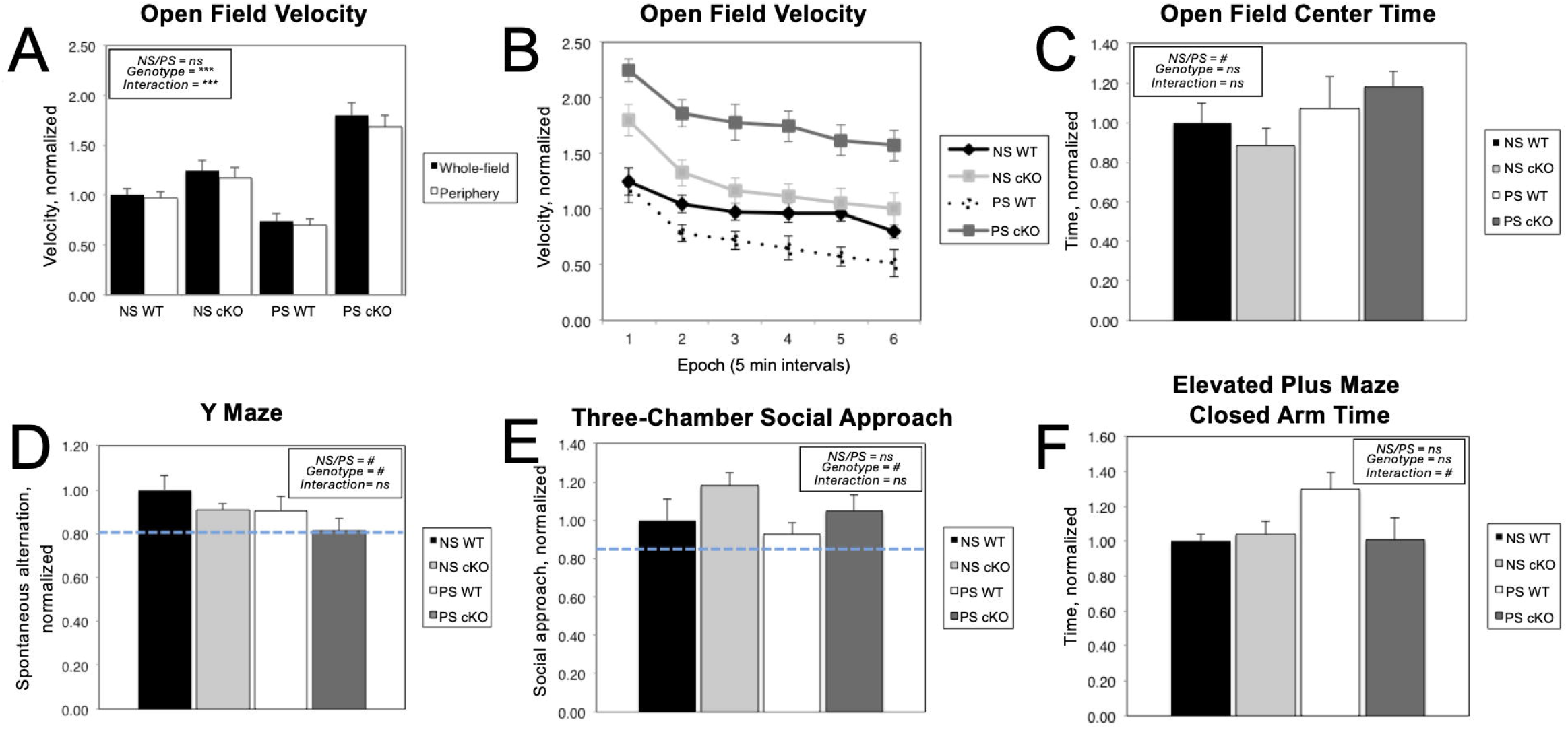
Prenatal stress may shift ADHD-relevant phenotypes of *Fgfr2* cKO males. All data are normalized to performance of the WT NS (control) group and presented as mean +/-standard error (SE). Two-way ANOVA results included in graphs: *\*\*\* = p<0*.*001; # = p < 0*.*1, ns= not significant*. **A)** Prenatal stress interacts with *Fgfr2* cKO to increase the whole-field velocity in the open field test, two-way ANOVA, *F*(1,47)=16.606, p < .0001. There is also a main effect of the Fgfr2 KO, *F*(1,47) = 42.576, p < .0001. n = 11 NS WT, 17 NS cKO, 12 PS WT, 11 PS cKO).**B)** Across-session normalized velocity per 5-minute epoch across the 30min open field-testing session supports group differences in average whole-field velocity findings (Figure 1A). **C)** Prenatal stress may result in impulsivity, measured as time spent in the center of the open field on the second day. There is a trend increase displayed by PS animals, *F*(1,46) = 2.990, p = .090. n = 11 NS WT, 17 NS cKO, 11 PS WT, 11 PS cKO mice. **D)** Prenatal stress may impair working memory, as measured by spontaneous alternation in the Y-maze. The dotted line represents the normalized chance level of alternation at 0.78. Trend decreases were observed in spontaneous alternation in PS offspring, *F*(1,48) = 3.154, p = .081 and cKO animals, *F*(1,48) = 2.848, p = .098. n = 13 NS WT, 16 NS cKO, 12 PS WT, 11 PS cKO mice. **E)** Social intrusiveness is not affected by prenatal stress, as measured by social approach in three chamber social task. The dotted line marks the normalized chance level of social approach at 0.83. The cKO groups display a trend increase in social approach, *F*(1,47) = 3.371, p = .073. n = 12 NS WT, 15 NS cKO, 13 PS WT, 11 PS cKO mice. **F)** Prenatal stress may protect against anxiety in a vulnerable genetic context, measured as time spent in the closed arm of the elevated plus maze. There is a trend interaction between prenatal stress and *Fgfr2* cKO to increase anxiety in the PS WT group, *F*(1,43) = 3.076, p = .087. n = 12 NS WT, 15 NS cKO, 9 PS WT, 12 PS cKO.

**Figure 3:**
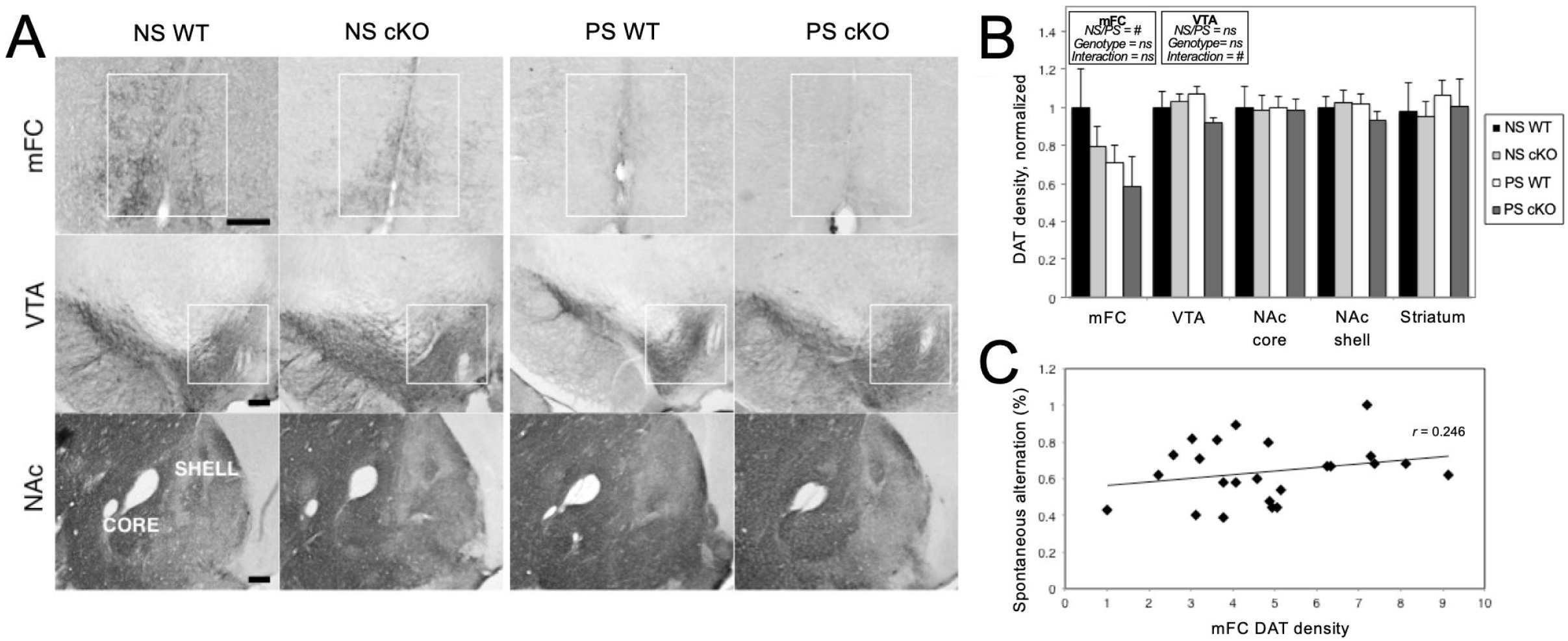
Prenatal stress may change DAT density in the mFC and midbrain VTA. Data normalized to NS WT (control) group and presented as mean +/-SE, unless otherwise noted. Two-way ANOVA results for mFC and VTA included in graphs: *# = p < 0*.*1, ns = not significant*. **A)** Representative images of staining for dopamine transporter (DAT; in dark) in the NS WT, NS cKO, PS WT and PS cKO groups, in the medial frontal cortex (mFC), ventral tegmental area (VTA), and nucleus accumbens (NAc), Scale bar is 100 μm. **B)** DAT density between groups. Trend decrease displayed by PS groups in the mFC, *F*(1,24) = 3.540, p = .072, and by PS cKO group in the VTA, *F*(1,24) = 3.767, p = .064. n = 5 NS WT, 9 NS cKO, 8 PS WT, 6 PS cKO. **C)** Correlation between spontaneous alternation and DAT density in the mFC, *r* = 0.246. Data used in correlations were not normalized.

PS offspring showed a trend decrease in DAT density in the medial frontal cortex (mFC) [*F*(1,24) = 3.540, *p* = .072], the projection area of DAergic neurons, exhibiting a 24% reduction relative to the NS animals (Figure 2B). Moreover, there was a trend interaction between prenatal stress and genotype in the ventral tegmental area (VTA) [*F*(1,24) = 3.767, *p* = .064]; there was a 13% decrease in PS cKO animals relative to PS WT and NS cKO groups. There were no significant effects in the nucleus accumbens (NAc) or the dorsal striatum. The pathway selectivity of these trend changes (e.g. primarily changes in the mesocorticolimbic (as opposed to the nigrostriatal) pathway) affected by prenatal stress is consistent with previous research (57, 61, 62).

At a group level, prenatal stress here altered ADHD-relevant behaviors and induced a trend for dopaminergic abnormalities. This is in accord with previous findings, but correlations between the two have not been examined previously. Therefore, potential correlations across individual animals were investigated between DAT density in regions affected by PS (the mFC and VTA) and behaviors for which prenatally stressed offspring exhibited ADHD-relevant characteristics (hyperactivity, impulsivity and impaired working memory) at trend or significant levels. In contrast to predictions, there were no significant or trend findings for any of the conducted correlations. To exemplify this: DAT density in the mFC plotted against spontaneous alternation is shown here (*r* = .246, N = 24, *p* = .248, two-tails) (Figure 2C).

## 4. DISCUSSION

Here, prenatal stress alone and in interaction with genetic susceptibility contributed to some ADHD-relevant outcomes. Previously, we found both embryonic dorsal forebrain and early postnatal astroglial, but not adult, deletion of *Fgfr2* (*Fgfr2* cKO) led to locomotor hyperactivity, among other ADHD-relevant behaviors (14). In the current study, prenatal stress further increased locomotor hyperactivity but did not specifically alter other behavioral phenotypes of *Fgfr2* cKO mice. Additionally, prenatal stress decreased DAT in the VTA only in *Fgfr2* cKO offspring. However, prenatal stress independently increased impulsivity, decreased working memory, and decreased medial frontal cortex DAT regardless of *Fgfr2* cKO effects on these outcomes. This supports prenatal stress as a contributor to ADHD-relevant outcomes, both in the presence and absence of genetic vulnerabilities.

Several genome-wide scans and association studies (GWAS) have attempted to understand the genetic etiology of ADHD (74-77), with the discovery of the first genome-wide significant risk loci in 2018 (76). These studies support the polygenic nature of ADHD, with more recent evidence supporting polygenic risk scores have relevance across the longitudinal course of ADHD and co-occurring psychiatric conditions (78). Interestingly, genetic variations of *Fgfr2* have generally not been associated with ADHD in GWAS studies. One genome-wide quantitative trait locus (QTL) study of Korean children with ADHD identified four QTLs, which did include a QTL with *Fgfr2* as a potential candidate (79). Rather, disruptions in the fibroblast growth factor system are increasingly linked with other psychiatric conditions such as anxiety and depression (80-83). Nonetheless, examining *Fgfr2* cKO and other models may provide valuable insight into neurobiological mechanisms of ADHD-relevant behavioral alterations. In the current study, we assessed DAT levels across ADHD-relevant brain regions, as disruption of DA systems is highly implicated in ADHD and the target of the most effective treatments. DAT impacts were not correlated with behavioral deficits. However, human GWAS and mouse transgenic models, including those manipulating FGF signaling, suggest multiple neuronal systems are involved in the pathophysiology of ADHD, including cortical interneurons (7, 9, 10, 84, 85) and synaptic development (14, 86, 87).

Prenatal stress has similarly been shown to affect such neuronal systems, including cortical and subcortical GABAergic cells: specifically, changes to the migration of interneuron progenitors, and reduced inhibitory functioning in the neonatal and adult forebrain have been shown (34). Prenatal stress has also been shown to impact development of dopaminergic systems (56, 88), consistent with the current study’s findings of DAT alternations in the medial prefrontal cortex.

Prenatal stress may alter dopaminergic system development, combining with genetic risks, through epigenetic mechanisms, as emerging evidence points toward prenatal stress leading to DNA methylation changes (89). Epigenetic changes may partially reflect gene-environment interactions underlying ADHD etiology. DNA methylation changes have also been found in both periphery and brain in individuals with ADHD (90-93). The difficulty of studying gene-environment interactions of ADHD is further complicated by the spectrum of individual differences in clinical presentation, rearing environments, and polygenic risk scores. One interpretation of the results here is that for ADHD, epidemiological studies of prenatal stress and GWAS studies may both fail to identify the contribution to risk if not accounting for effects from the other variable. It is a challenge to account for the most important additional risk variables that may have interactive impacts on association studies. A study exploring the interaction between prenatal stress and polygenic risk scores for ADHD found prenatal stress may promote slower adolescent neurodevelopment in individuals with higher ADHD polygenic risk scores, but promote faster brain development in individuals with lower ADHD genetic vulnerability (94).

Gene-environment interactions are strongly implicated in the differential-susceptibility framework (95). While individual differences in responses to environmental factors have been well-appreciated, this framework counters the narrative that “vulnerable” individuals are disproportionately affected by *only* adverse experiences. Rather, individuals carrying different genetic markers may have differential susceptibility to *both* positive and negative experiences. In the current study, prenatally stressed WT animals displayed increased anxiety-relevant behavior, but prenatally stressed *Fgfr2* cKO mice did not show this increase. This finding may suggest potential protective factors from genetic “vulnerabilities” for certain behavioral phenotypes. This framework also highlights opportunities for interventional strategies that include minimizing adverse experiences as well as increasing positive experiences in modifying childhood psychiatric disease risk. Rodent studies of neurodevelopmental disorders models show enriched environments ameliorate behavioral abnormalities of transgenic mice, including social interaction and working memory deficits and hyperactivity in Fragile X syndrome mouse models (96, 97). These findings from enriched environment studies may be particularly relevant to ADHD. Evidence suggests children with genetic variants of dopamine receptor D4 (an identified vulnerability factor to ADHD (98)) are not only more adversely affected by poorer quality parenting, but they also benefit more from higher quality environments (95, 98). While positive rearing environment models have not yet been tested in *Fgfr2* cKO mice and no postnatal manipulations were done in the current study, postnatal rearing environments will be an important future direction to understand how they modify risk from ADHD-relevant genetic vulnerability and gestational stressors.

Overall, these data support prenatal stress as a contributor to ADHD-relevant outcomes, both independently and with genetic vulnerability from a *Fgfr2* cKO mouse model. Findings from this study aid in our broader understanding of how environmental stressor impacts on embryonic brain development affect long-term behavioral and neurobiological outcomes, as well as how prenatal environments may modify risk from genetics. Improving our understanding of environmental modifiers of genetic risks for common childhood psychiatric conditions will be beneficial for designing intervention strategies to minimize the burden of these disorders across the lifespan.

## ACKNOWLEDGEMENTS

The authors would like to thank Dr. Flora Vaccarino for generously providing mice and laboratory space, and for helpful discussion during the course of the study. The authors would like to acknowledge the following funding sources: NARSAD/Brain Behavior Research Foundation Young Investigator Award (HES), NINDS grant T32NS007124 (MXC), NIMH grant K08MH086812 (HES), and the University of Iowa Ida P Haller Chair in Child and Adolescent Psychiatry (HES). Figure 1 was created using BioRender.com.

## DISCLOSURES

The authors declare they have no conflicts of interest.

## Notes

### Competing Interest Statement

The authors have declared no competing interest.

